# Computational Simulations of Hyoid Bone Position and Tracheal Displacement: Effects on Upper Airway Patency and Tissue Mechanics

**DOI:** 10.1101/2024.08.19.608628

**Authors:** Dana Bekdache, Jason Amatoury

## Abstract

Surgical hyoid repositioning (HR) improves upper airway (UA) patency. Tracheal displacement (TD) is likely to impact HR outcomes, and vice versa, due to hyoid-trachea connections. This study used computational modeling to investigate the influence of TD and HR on UA outcomes and examine the impact of a more caudal baseline hyoid position (OSA phenotype).

**Methods:** A 2D finite element model of the rabbit UA was used to simulate TD and HR (in different directions), separately and combined. Model outcomes included UA closing pressure (Pclose), area, anteroposterior diameter (APD) and soft tissue mechanics (stress/strain). Simulations were repeated with a more caudal baseline hyoid position.

**Results:** Compared to baseline (TD=HR=0mm), TD alone reduced Pclose by −34%, increased area by 21% and APD by up to 18%. HR alone (except caudal) improved outcomes, particularly anterior-cranial HR which decreased Pclose by −106%, increased area by 32% and APD by up to 107%. TD+HR (except caudal) enhanced these outcomes, with TD+anterior-cranial HR further decreasing Pclose (−131%) and increasing area (55%) and APD (128%). A more caudal baseline hyoid position reduced the effect of TD+anterior-cranial HR on Pclose (−43%), area (49%) and APD (115%).

**Conclusion:** The combination of TD and HR (except caudal) improved UA outcomes even further than when either intervention was applied alone. A more caudal baseline hyoid position reduced the overall impact of each intervention. This study suggests that considering the baseline hyoid position, the degree of TD, and the extent/direction of surgical HR could be crucial in optimizing OSA treatment outcomes.

**Key points summary:**

- Surgical hyoid repositioning can improve upper airway patency and is a treatment for obstructive sleep apnea (OSA).
- Tracheal displacement, also critical to upper airway function, likely influences hyoid repositioning outcomes due to hyoid-trachea connections.
- This study used a computational model of the upper airway to simulate tracheal displacement and hyoid repositioning in various directions and magnitude, assessing their impact on upper airway collapsibility, size, and soft tissue mechanics. The influence of a more caudal baseline hyoid position, like in OSA, was also simulated.
- Combining tracheal displacement with anterior-based hyoid repositioning, in particular, resulted in greater improvements in upper airway outcomes compared to tracheal displacement and hyoid repositioning alone.
- A more caudal baseline hyoid position diminished the upper airway improvements with both interventions
- Optimizing OSA treatment outcomes with hyoid surgeries may require considering the baseline hyoid position, the degree of tracheal displacement, and the direction/magnitude of surgical hyoid repositioning.

## INTRODUCTION

Obstructive sleep apnea (OSA) is a disorder where the upper airway collapses repeatedly during sleep and obstructs airflow to the lungs despite the presence of respiratory effort. OSA is a highly prevalent disorder affecting nearly 1 billion people worldwide (1) and is linked with health consequences including excessive daytime sleepiness, reduced quality of life, cardiovascular anomalies and neurocognitive disorders (2, 3).

One anatomical factor that influences the upper airway is the hyoid bone, which is a floating and mobile bone at the midline of the neck, sitting just below the base of the tongue. It has muscle and other anatomical connections to the thyroid cartilage, pharynx, tongue and mandible. As a result of the hyoid bone’s diverse connections and strategic position, it plays an essential role in upper airway function such as breathing, speaking, swallowing and chewing (4–6). The hyoid is also connected to several upper airway dilator muscles in the neck. Accordingly, the hyoid’s position is important in supporting pharyngeal muscle passive and active function for maintaining upper airway patency (4, 7). A common anatomical phenotype amongst individuals with OSA is a more inferiorly (caudally) positioned hyoid bone, resulting in a more collapsible upper airway (8, 9). Surgical hyoid bone repositioning (HR) has been a viable treatment option to improve upper airway patency in OSA, yet it is unreliable, with success rates ranging between 17-76% (10–13).

Another key factor in maintaining upper airway patency is the trachea. During inspiration, an increase in negative intraluminal pressure and lung volume occurs, resulting in caudal tracheal displacement (TD). This in turn, enlarges the upper airway, stiffens its tissues and decreases pharyngeal tissue pressure (14–17). Apart from this dynamic displacement, the trachea also experiences static displacement from different sleeping positions and weight changes (18, 19). Additionally, TD also results in hyoid displacement due to anatomical connections (15). However, the influence of hyoid bone position and TD on each other and upper airway patency has yet to be explored.

Therefore, given the important role of both the hyoid and the trachea in maintaining upper airway patency and their anatomical connection, it is essential to understand how TD affects HR surgery outcomes (and vice versa). Furthermore, given the lower position of the hyoid in OSA, investigating the effect of baseline hyoid position (phenotype) on the outcomes of both TD and HR is also of particular interest.

In-vivo research in animals regarding the hyoid and the trachea is invasive and challenging due to the complexity of the upper airway and the difficulty to control all physiological and anatomical factors at once. Furthermore, altering the natural baseline hyoid bone position is not possible using experimental models. Accordingly, computational modeling is required. We have previously developed and validated a computational model of a rabbit upper airway that allows for such studies to be undertaken (16, 20). The rabbit, with its anatomical resemblance to humans, has consistently demonstrated similarities in upper airway outcomes to the human (4, 15–17, 21–24), so it is an ideal model to study the upper airway.

The current study aimed to utilize our finite element model to simulate TD and surgical HR, separately and in combination, to assess their effects on upper airway collapsibility, size and soft tissue mechanics. Moreover, this work explores how a more caudal baseline hyoid bone position (OSA phenotype) influences TD and surgical HR upper airway outcomes.

## METHODS

A validated mid-sagittal 2D computational finite element model of the passive rabbit upper airway was used to undertake simulations. The model was originally developed and validated by Amatoury et al (16) to allow for TD and mandibular advancement simulations. It was then subsequently advanced and independently validated by Salman and Amatoury (20) to allow for changes in baseline hyoid position (phenotype) and surgical HR, and application of intraluminal upper airway pressure (Pua) for quantification of upper airway collapsibility. The model of Salman and Amatoury (20) was utilized in this study to undertake simulations of TD, surgical HR, changes in baseline hyoid position and Pua. The model has been described in detail previously (16, 20) and only a summary is provided below with aspects associated with this study explained in greater detail.

### Model Geometry and Mesh

The model geometry was developed from a CT image of the rabbit upper airway (Figure 1A) and includes the mandible, hard palate, hyoid bone, thyroid cartilage, epiglottis, tongue, soft palate, constrictor muscles (represented as a single body), geniohyoid and mylohyoid muscles, in addition to a soft tissue mass (Figure 1B).

**Figure 1:**
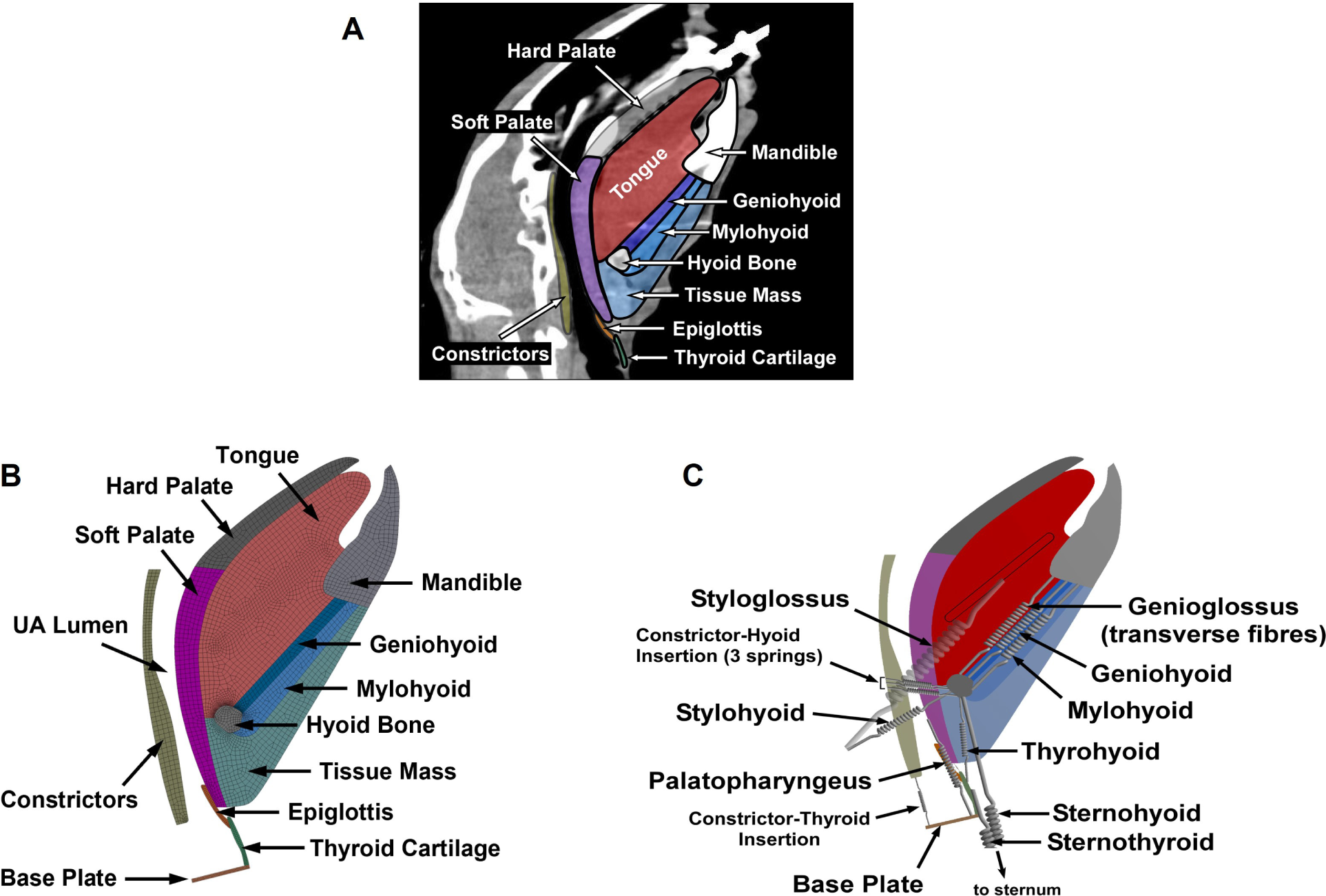
Model of the Rabbit Upper Airway. (A) Midsagittal CT image of a rabbit upper airway used for developing the 2D FE model geometry. The colored regions represent the tissues represented in the model. (B) The FE model reconstructed from the CT image. (C) Muscles represented as linear elastic springs to mimic their function and capabilities (spring properties are summarized in Figure S3B). Figure adapted and modified from Salman and Amatoury (20).

A 2D mesh composed of four-node quadrilateral-dominant plain-strain elements was created from the geometry. The resulting mesh consisted of a total of 3,970 elements, comprising 3,907 quadrilateral elements and 63 triangular elements (Figure 1B).

### Material Properties, Boundary & Contact Conditions

The linear elastic material properties of the boney and cartilaginous tissues and hyperelastic soft tissue components of the model are summarized in Supplementary Material Tables S1 and S2.

Model boundary and contact conditions are summarized in Supplementary Material Figures S1 and S2, respectively. The passive action of several upper airway muscles was represented as springs (Figure 1C), and their properties are summarized in Figure S3.

### Loading Conditions and Simulation Protocol

Initially, TD was simulated by applying a caudal displacement load perpendicular to the thyroid cartilage via the base plate, which the trachea directly connects to, as described previously (16) (Figure 2). TD was applied in increments of 1 mm from 0-3 mm. Note that this displacement translates to 0, 1.61, 3.23, and 4.84 mm of TD applied between the 3^rd^/4^th^ cartilaginous rings experimentally in rabbits (15). HR was then applied alone as a displacement vector in increments of 1 mm from 0-3 mm in 5 different directions: anterior, cranial, caudal, anterior-cranial (45°) and anterior-caudal (45°). Following this, for each HR direction/increment, TD was applied in increments of 1 mm from 1-3 mm. After each displacement application (HR alone, TD alone and the combination of both TD and HR (TD+HR), a negative upper airway intraluminal pressure (Pua) of up to an equivalent of ∼ −8 cm H_2_O was applied on the walls of the upper airway, as per Salman and Amatoury (20) (Figure 2), to quantify closing pressure (see below).

**Figure 2:**
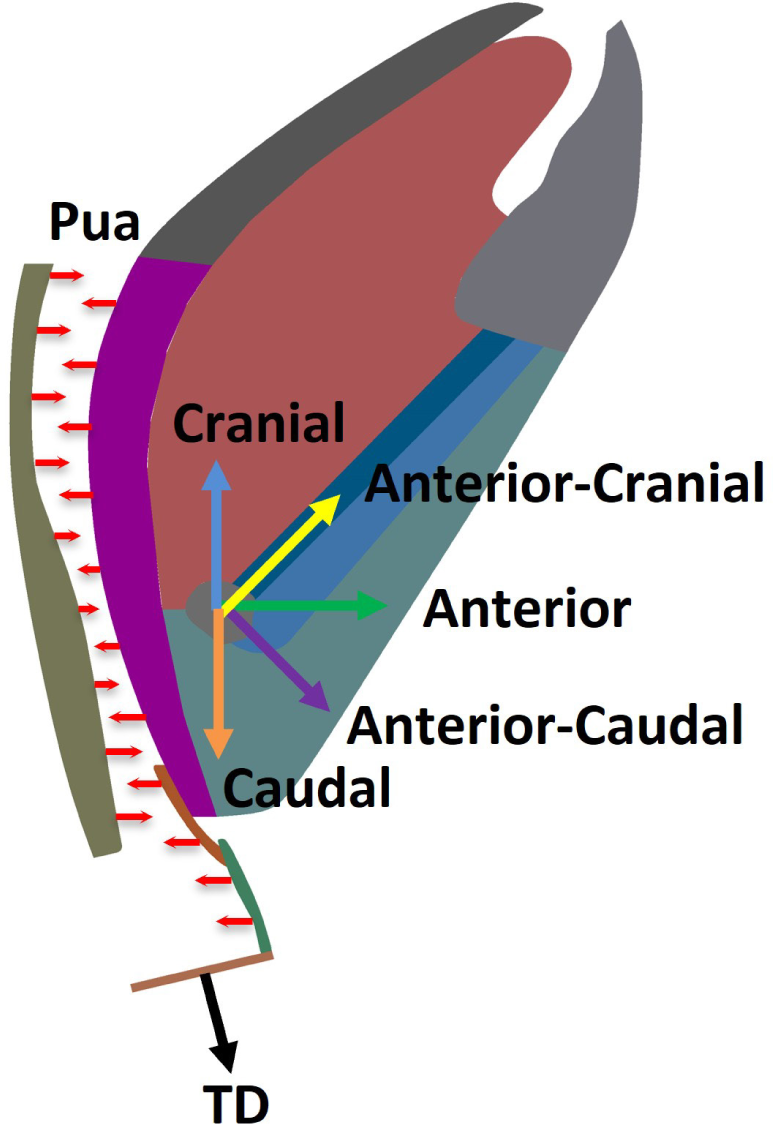
Loading Conditions. Tracheal displacement (TD) was applied perpendicular to the thyroid cartilage base plate. Surgical hyoid repositioning (HR) was applied in 5 different directions (as shown). Negative intraluminal pressure (Pua) was applied on the upper airway luminal walls for quantification of closing pressure (Pclose). Springs (passive muscles) have been omitted for improved visualization.

Simulations were also undertaken with the baseline hyoid position placed in a more caudal location, mimicking the OSA phenotype. This was achieved by shifting the hyoid’s baseline position geometrically by −2 mm and −4 mm in the caudal direction (Figure 3). For each caudal hyoid positioned baseline geometry, the clinically relevant anterior-cranial HR, akin to hyomandibular suspension, was applied alone and in combination with TD, followed by Pua.

**Figure 3:**
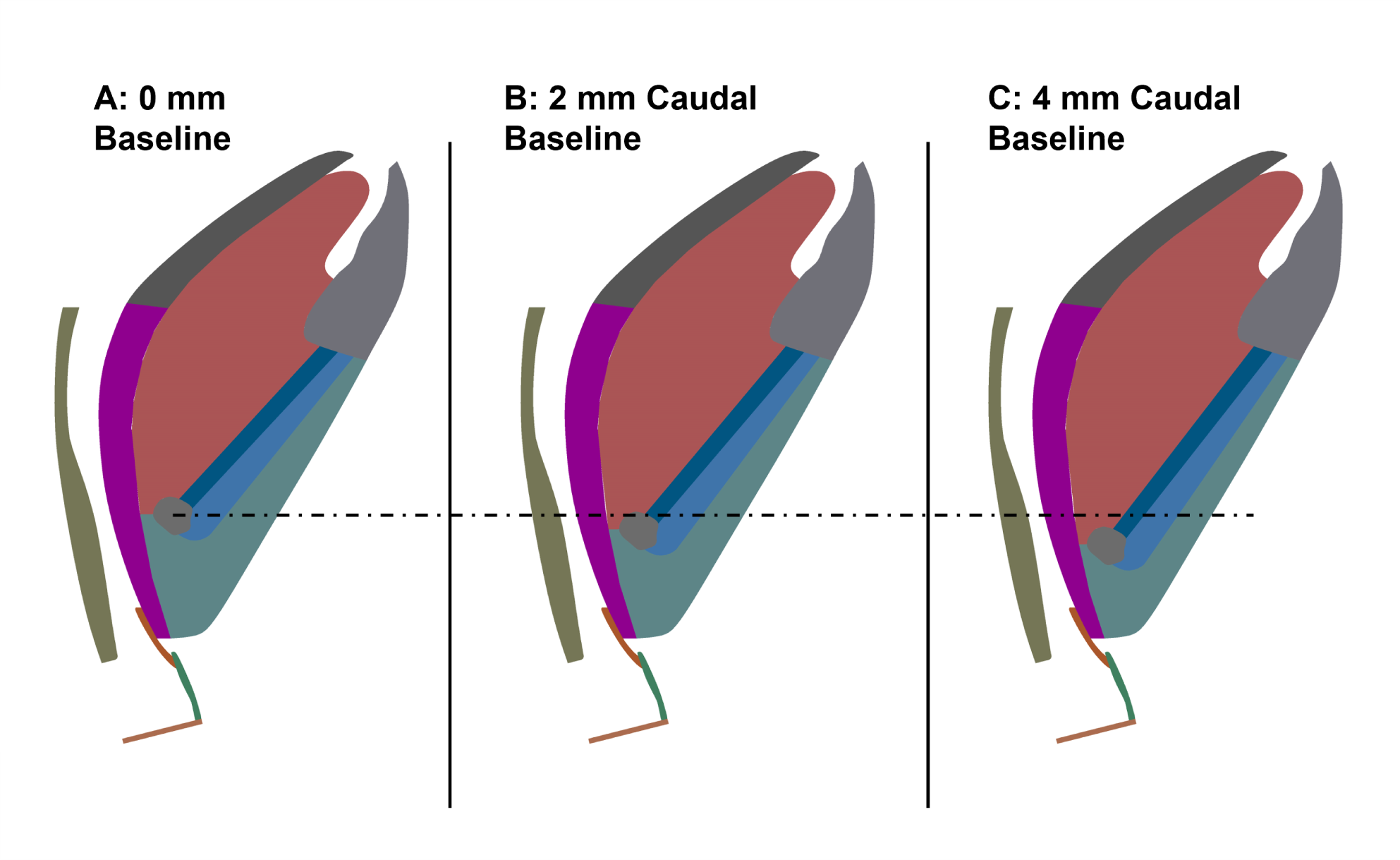
Baseline Hyoid Positions. The baseline hyoid position was shifted caudally 2 mm (B) and 4 mm (C) to represent the OSA phenotype and then select simulations were undertaken with this new geometry. The dotted black line represents the centroid level of the hyoid bone in its original (0 mm) baseline position. Springs have been omitted for improved visualization.

### Finite Element Analysis Settings

Finite element analysis was performed using ANSYS Workbench using a static structural component system (release 19.2; ANSYS Academic Research, Canonsburg, PA). The system was applied with 2D plane strain conditions and large deformation effects since the model consists of non-linear hyper-elastic materials with expected large deformations (16, 20). TD and HR were performed individually over one step, followed by Pua as an additional step, using 80 equally spaced substeps (see *Upper Airway Collapsibility* below) (20). For TD+HR, HR was performed in the first step, TD in the second step and then Pua in the third step.

### Model Outputs

#### Upper Airway Collapsibility (Closing Pressure; Pclose)

Upper airway collapsibility was quantified using closing pressure (Pclose), the pressure at which the upper airway closed. Following application of Pua, the substep at which upper airway closure occurred was recorded using the ANSYS contact tool (20). Pclose was then calculated and expressed as a percent change relative to the original baseline hyoid position (HR=TD=0mm; ΔPclose).

For certain HR directions at high TD and HR increments where the model failed to converge to collapse with the Pua load (only 3 instances), a liner regression model was fitted to existing data and the missing points then estimated using the model (GraphPad Prism version 9.1.1; GraphPad Software Inc., La Jolla, CA).

#### Upper Airway Lumen Geometry

For each independent or combined TD and HR increment, the deformed geometry was exported from ANSYS as a STEP file into Rhinoceros V5 (Robert McNeel and Associates, Seattle, WA) to extract the upper airway lumen for geometry analysis. A midsagittal contour of the upper airway was created (Figure 4) with the upper (cranial) boundary being the nasal choanae level and the lower (caudal) boundary being the base of the epiglottis (Figure 4A). The geometry was analyzed for mid-sagittal cross-sectional area (Area) and anteroposterior diameter (APD) (Figure 4B). APD was obtained along the length of the upper airway every 0.5 mm from the cranial boundary and then averaged across three regions (16). These regions included (Figure 4A): R1, from the nasal choanae level to the upper surface of the hyoid bone; R2, from the base of R1 to 1.5 mm above the upper epiglottis surface; R3, from the base of R2 to the base of the epiglottis (surface of the glottis). Area and average region APDs were expressed as a percent change relative to their respective baselines (ΔArea and ΔAPD, respectively).

**Figure 4:**
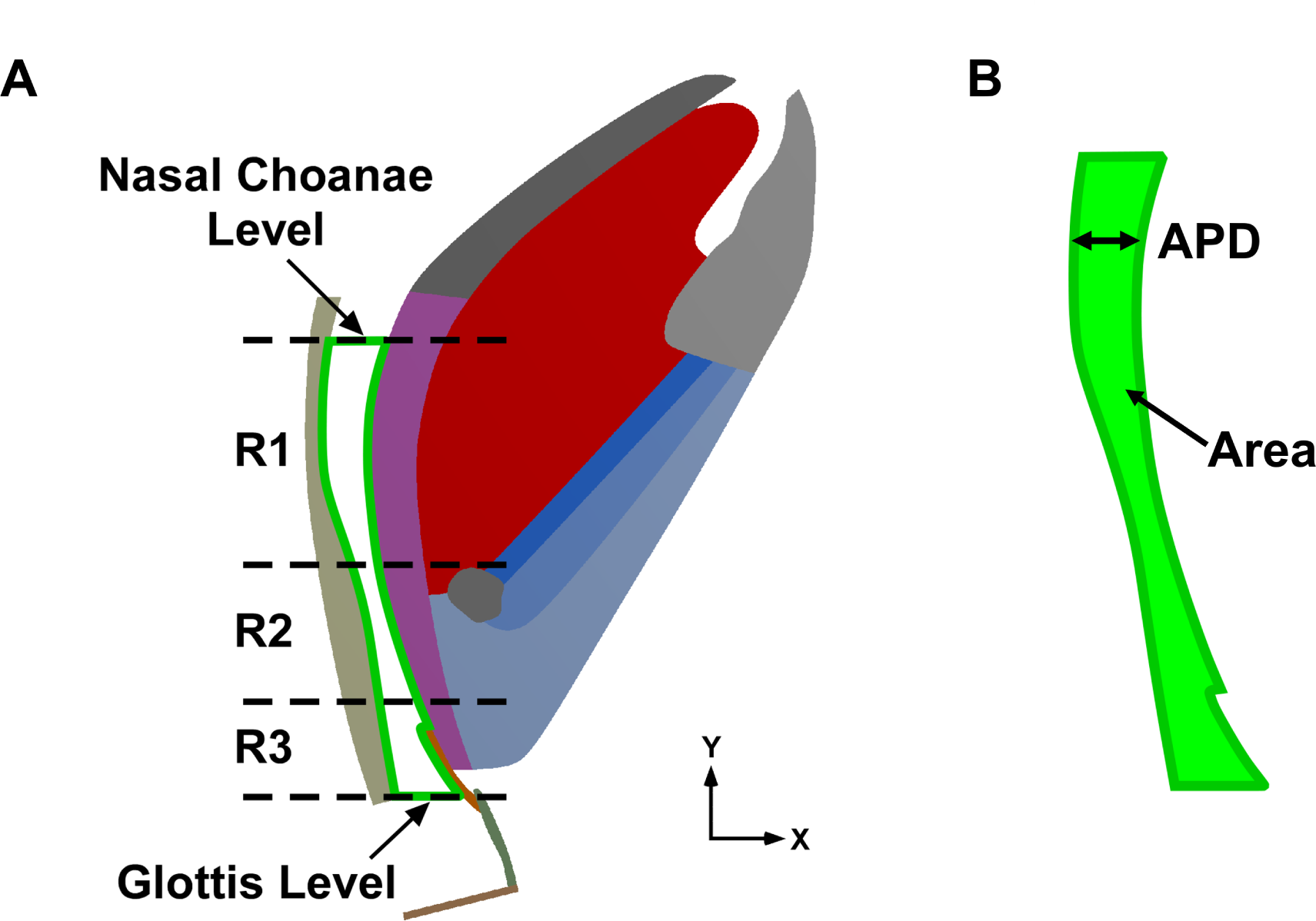
Upper Airway Lumen Geometry. (A) Upper airway lumen (outlined in green) for the undeformed upper airway model, with nasal choanae and glottis levels indicated. Upper airway regions are also shown, where R1 extends from the nasal choanae level to the upper surface of the hyoid bone, R2 from the hyoid bone to 1.5 mm superior to the epiglottis surface, and R3 from the base of R2 to the base of the epiglottis (surface of the glottis). Springs have been omitted for improved visualization. Adapted from Amatoury et al (16). (B) The upper airway segment extracted and assessed for its cross-sectional area (Area), highlighted in light green, and the anteroposterior diameter (APD), measured from the constrictor to the soft palate as indicated. APD measurements were taken 0.5 mm apart from the nasal choanae level and averaged for each region.

#### Upper Airway Soft Tissue Mechanics (tissue displacement, stress and strain)

The resultant displacement of all upper airway tissues was obtained and presented by color-coded contour maps (resultant vectors) of the deformed meshed model. Similarly, von Mises stresses and strains of the soft tissues were also obtained.

## RESULTS

### Surgical Hyoid Repositioning and Tracheal Displacement

Here, results are presented for TD and HR simulations when the hyoid bone was at the original baseline position.

#### Upper Airway Collapsibility (Pclose)

Figure 5 summarizes Pclose outcomes with TD and HR. TD alone resulted in a progressively more negative Pclose compared to baseline (less collapsible), with a reduction of up to −34% at 3 mm. Similarly, HR alone (except caudal) led to a more negative Pclose with each HR increment, with the anterior-cranial direction demonstrating the greatest change of −106% at 3 mm, while the least decrease occurred in the cranial direction with a change of −26% at 3 mm. The caudal HR direction had the opposite effect, with up to a 20% increase in Pclose. TD+HR resulted in the greatest decrease in Pclose compared to TD or HR alone, with anterior-based HR directions causing the greatest decreases, particularly anterior-cranial with Pclose going down to −131% at 3 mm and cranial down to −66%. For TD+caudal HR, a maximum Pclose change of −6% was observed (Figure 5).

**Figure 5:**
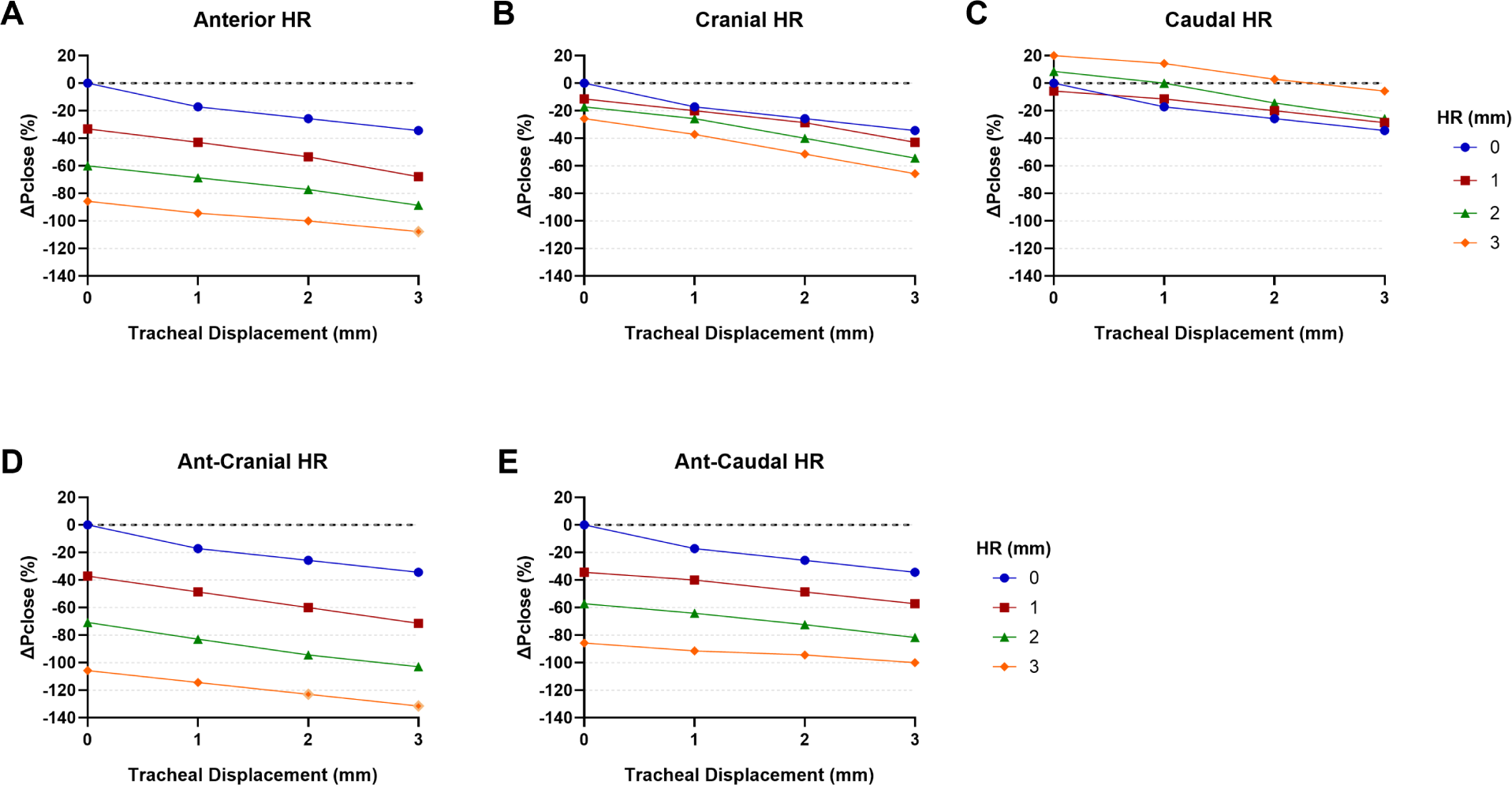
Pclose Outcomes for the Original Baseline Hyoid Position. Percent change in Pclose (for to the original baseline hyoid position) with tracheal displacement (TD) and hyoid repositioning (HR) in each direction (A-E). Increasing TD alone (HR=0mm; blue solid circles) progressively decreased Pclose (A-E). Increasing HR alone (TD=0mm; different symbols/colors intersecting the y-axis) also progressively decreased Pclose in all directions, except for caudal HR (C) that demonstrated a slight increase. Anterior-based HR directions demonstrated the greatest change in Pclose (A, D, E), particularly anterior-cranial (ant-cranial) HR. TD+HR decreased Pclose even further then when either was applied alone, and this effect was near additive. Caudal HR negative the decreased in Pclose with TD (C). Note that for TD=3 mm + ant-cranial HR=3 mm (A), TD=2 mm + ant-cranial HR=3 mm and TD=3 mm + ant-cranial HR=3 mm (D), open diamonds represent points obtained using linear regression.

#### Upper Airway Geometry

##### Area

Figure 6 summarizes Pclose outcomes with TD and HR. TD alone increased upper airway area by up to 21% at 3 mm. HR alone increased area in all directions, except for caudal HR. Anterior-cranial HR had the greatest increase of 32%, while cranial HR only slightly increased area by 4%, which was less than the increase due to TD alone. Conversely, caudal HR alone resulted in a slight decrease in area of −5%. TD+HR resulted in a greater increase in area compared to TD or HR alone, particularly for anterior-based directions and mostly with TD+anterior-cranial HR with area now going up by 55% at 3mm. However, TD+caudal HR resulted in a lower change in area (16%) compared to the application of TD alone (21%) (Figure 6).

**Figure 6:**
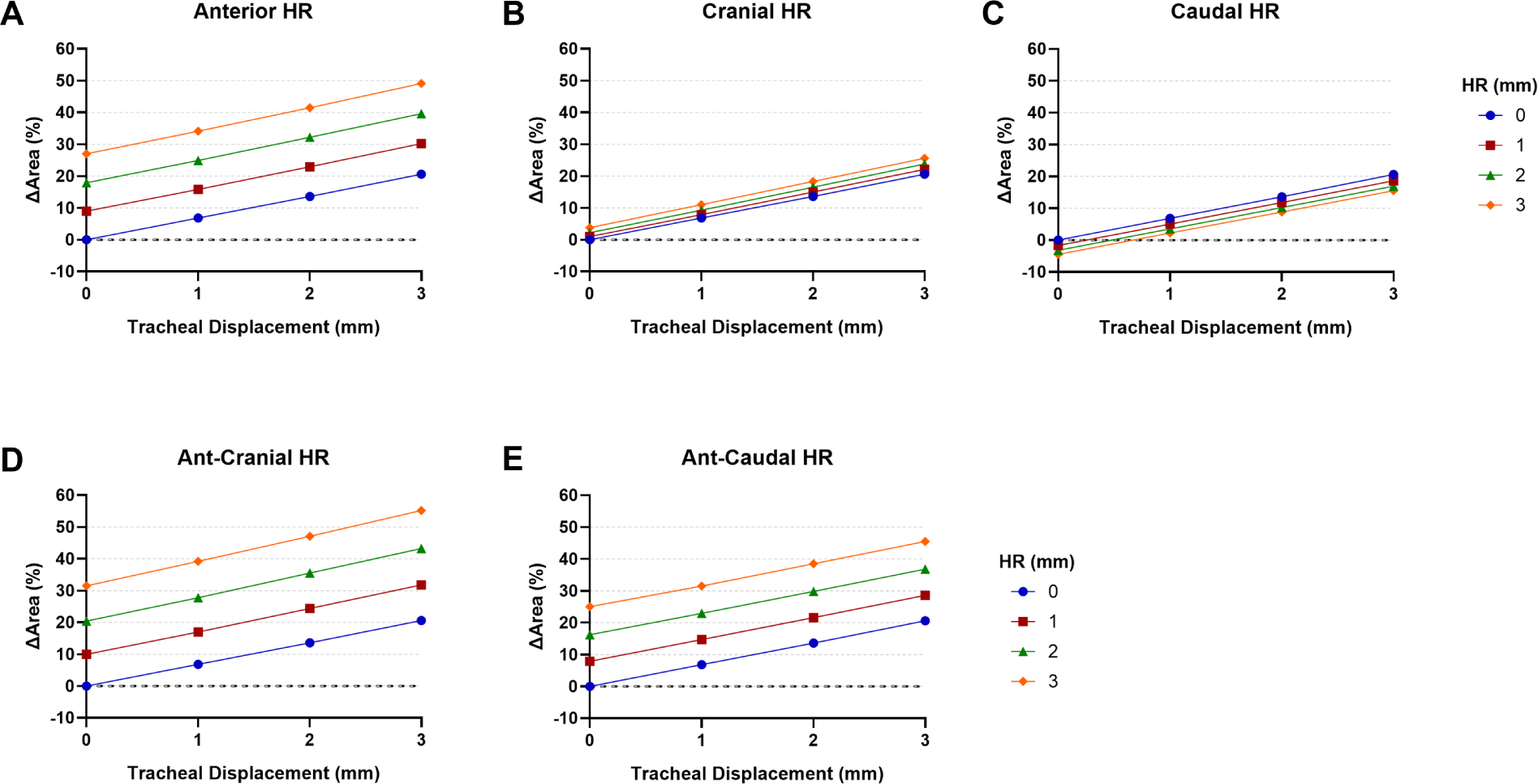
Upper Airway Area Outcomes for the Original Baseline Hyoid Position. Percent change in Area (ΔArea) with tracheal displacement (TD) and hyoid repositioning (HR) in each direction compared to baseline for the original baseline hyoid bone position. Area increased with increasing increments of TD or HR alone, except caudal HR which demonstrated a decrease. Anterior-based HR directions exhibited the greatest change in Area, and mostly anterior-cranial (ant-cranial) HR. The combination of TD and HR resulted in even greater increases in Area that TD or HR alone, in an almost near additive manner. The anterior-based directions demonstrated the greatest increase in area with TD, and most notably TD+ant-cranial HR.

##### Anteroposterior Diameter

Figure 7 summarizes APD changes with TD and HR. TD alone increased APD for all regions, with the highest change occurring in R2 (18% at 3mm). HR alone also increased APD compared to baseline for all regions, except for R1 in cranial (−12% at 3 mm), R2 (−25% at 3 mm) and R3 (−15% at 3 mm) in caudal, and R3 in anterior-caudal (−5% at 3 mm). Anterior-cranial HR resulted in the greatest increase in R2 (107% at 3mm).

**Figure 7:**
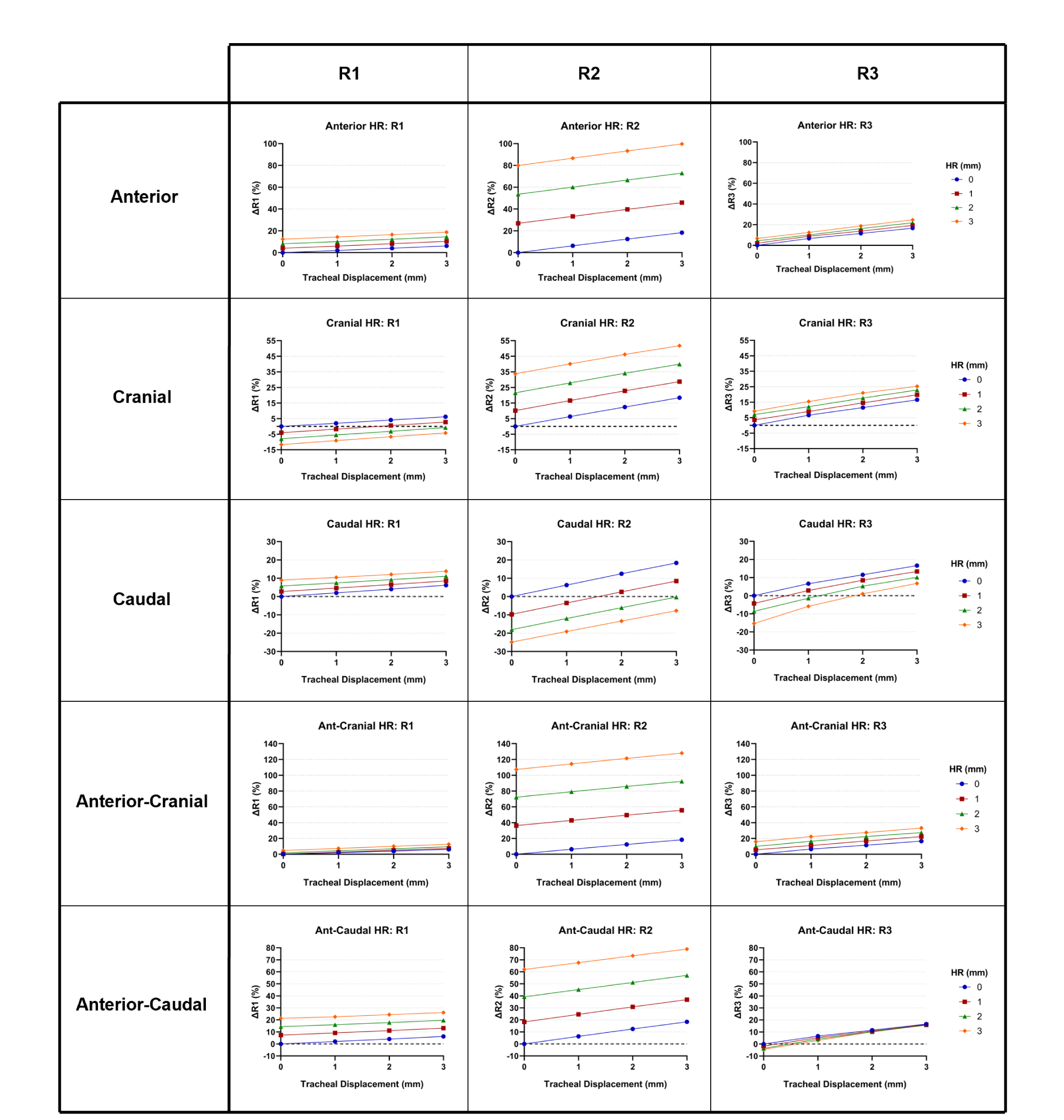
Upper Airway APD Outcomes for the Original Baseline Hyoid Position. Percent change in anteroposterior diameter (APD) with tracheal disaplcement (TD) and hyoid repositioning (HR) in each upper airway region (R1, R2 and R3). Increasing TD or HR alone resulted in an increase in APD in all upper airway regions, especially in region R2, except for caudal HR that showed a decrease in R2 and R3 when caudal HR is increased. When TD and HR were combined, APD increased even further than when TD or HR were applied alone, particularly for anterior-based directions, and mostly anterior-cranial (ant-cranial) HR. The largest changes in APD for TD+HR occurred in R2.

TD+HR together resulted in the greatest increase in APD for all regions, with the anterior-cranial direction showing the largest change. The R2 region showed the highest change in APD across all HR directions (except caudal), with the TD+anterior-cranial HR going up to 128%. Conversely, the TD+caudal HR change in APD of R2 was negative (−8%), yet it was an improvement from the R2 of caudal HR alone (−25%).

#### Upper Airway Soft Tissue Mechanics

Figures 8-10 show contour maps of stress, strain and displacement. TD alone increased stress and strain, particularly along the constrictors (Figure 8 and 9). HR alone also increased stress and strain. TD+HR resulted in a greater increase in stress and strain (except caudal HR), particularly for the anterior-based directions, and mostly anterior-cranial. TD+caudal HR resulted in lower stress and strain around the hyoid bone than when HR was applied alone (Figures 8 & 9).

**Figure 8:**
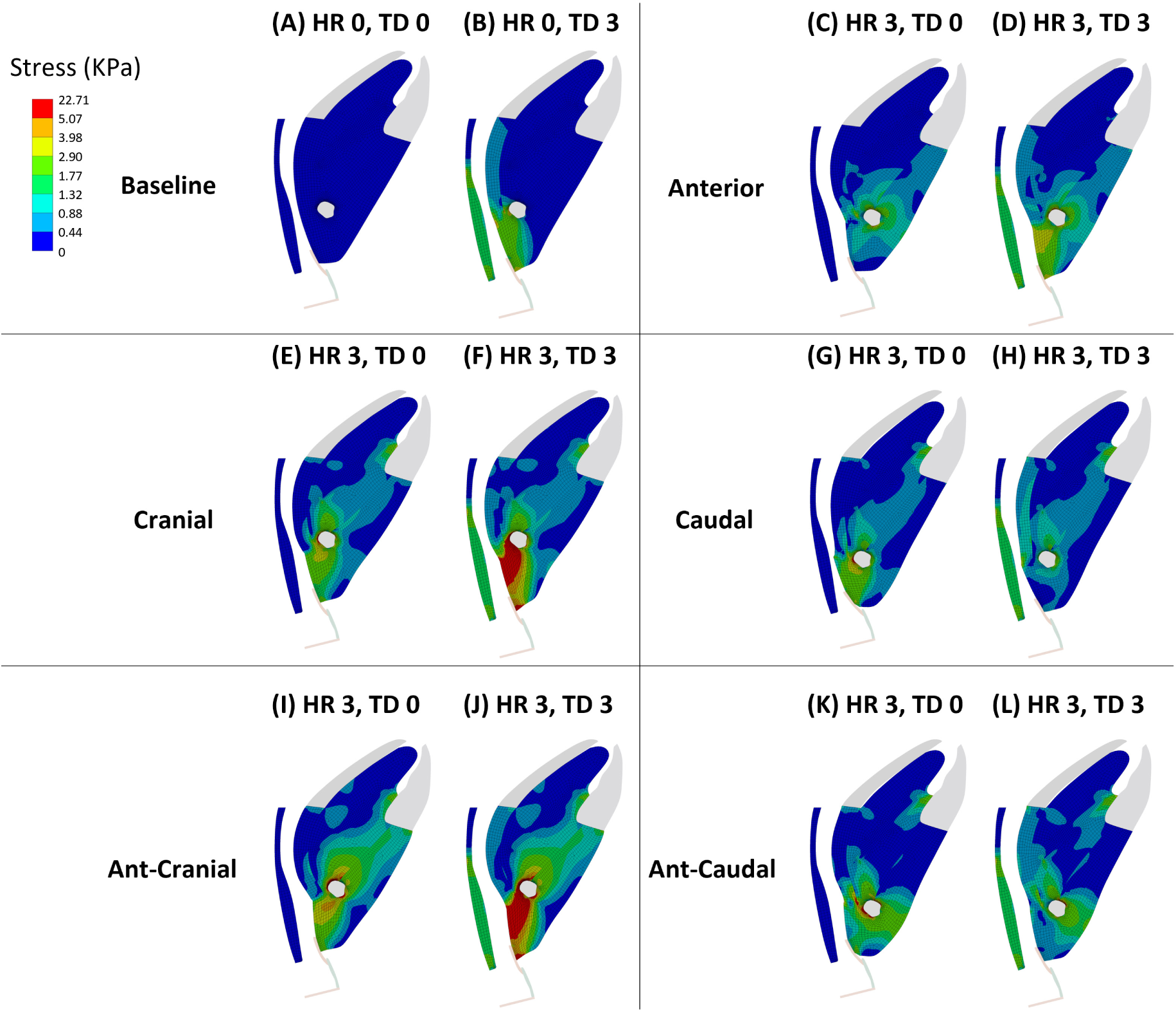
Soft Tissue Stress for the Original Baseline Hyoid Position. Tracheal displacement (TD) and hyoid repositioning (HR) alone both resulted in increased stress distributions along different soft tissue regions. TD alone resulted in stresses concentrated along the soft palate and constrictor muscles (B). HR alone increased stresses throughout different soft tissue regions depending on the HR direction. TD+HR resulted in the greatest distribution of stresses, especially for anterior-cranial (ant-cranial) HR (J). However, TD+caudal HR resulted in a lower stress distribution around the hyoid bone than when caudal HR was applied alone. HR 0 = HR at 0 mm. HR 3 = HR at 3 mm. TD 0 = TD at 0 mm. TD 3 = TD at 3mm.

**Figure 9:**
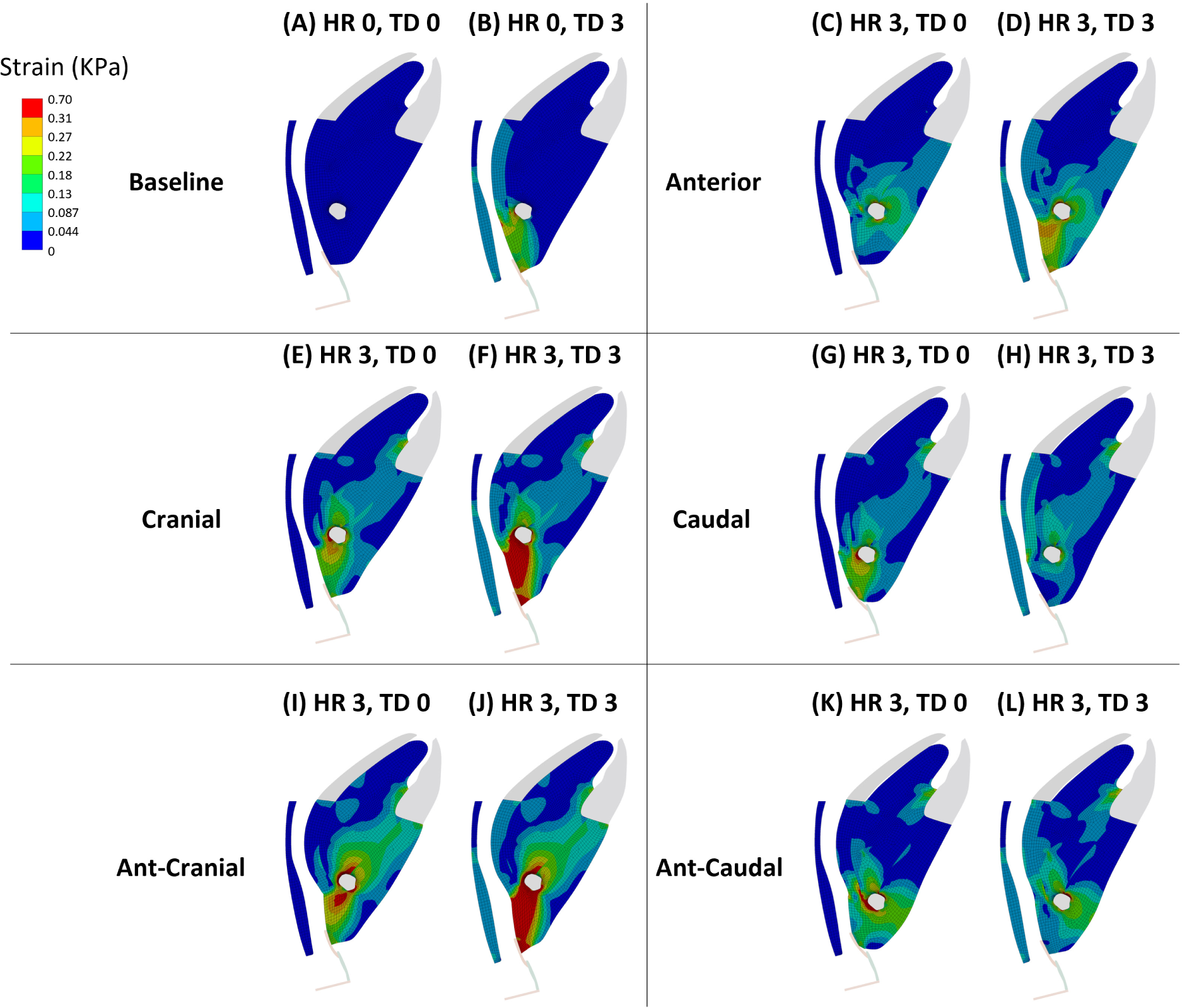
Soft Tissue Strain for the Original Baseline Hyoid Position. Tracheal displacement (TD) and hyoid repositioning (HR) alone resulted in different strain distributions along the soft tissues. However, TD+HR resulted in the greatest distribution of strains especially for anterior-cranial HR (J). However, TD+caudal HR resulted in a lower strain distribution around the hyoid bone than when HR was applied alone. HR 0 = HR at 0 mm. HR 3 = HR at 3 mm. TD 0 = TD at 0 mm. TD 3 = TD at 3mm.

**Figure 10:**
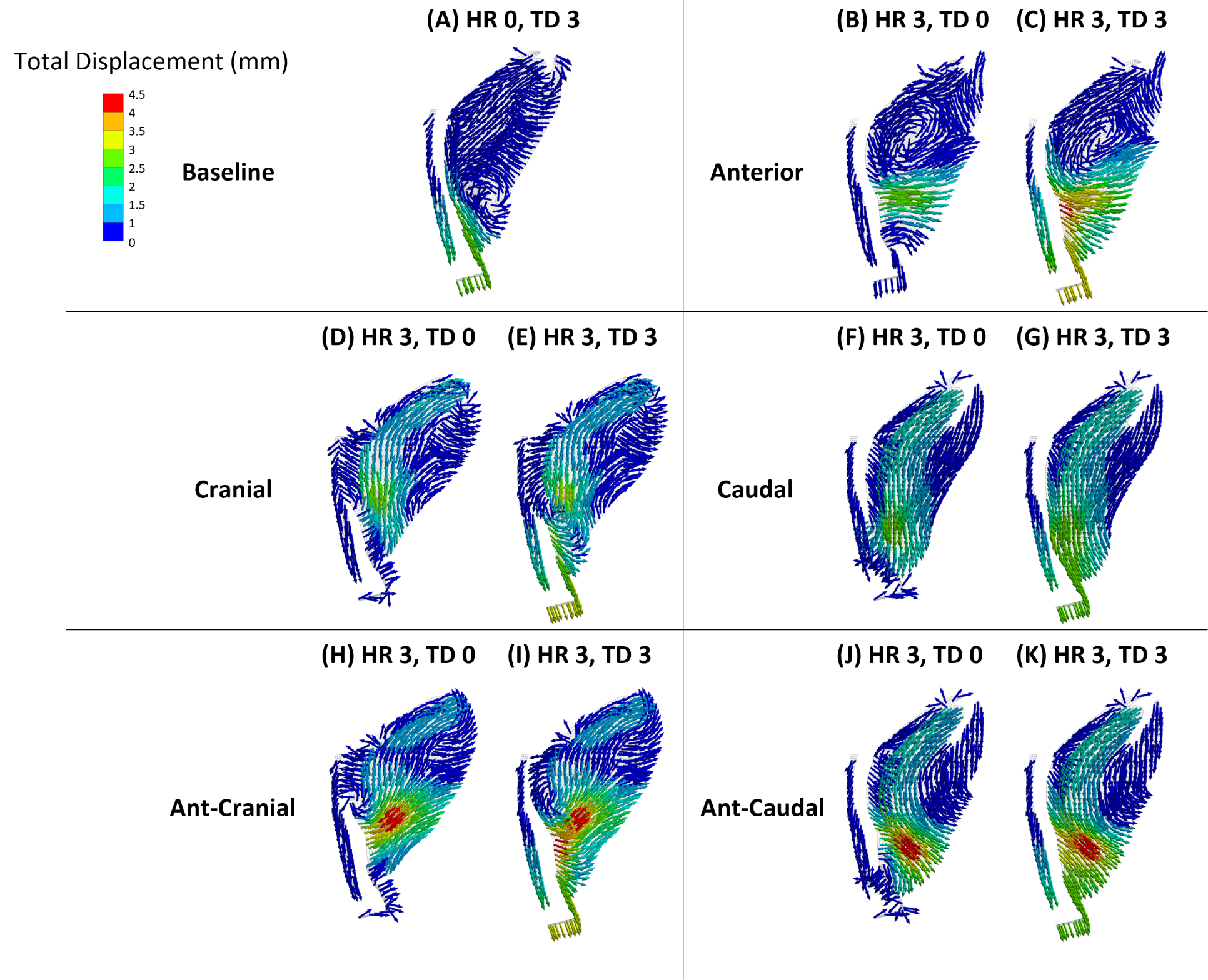
Tissue Displacement for the Original Baseline Hyoid Position. The total (resultant) displacement contour maps indicate that tracheal displacement (TD) and hyoid repositioning (HR) alone resulted in an overall increased displacement distribution in the indicated vector directions. The combination of TD and HR resulted in greater tissue displacements, especially with anterior-cranial HR (I). HR 0 = HR at 0 mm. HR 3 = HR at 3 mm. TD 0 = TD at 0 mm. TD 3 = TD at 3mm.

The contour maps for stress and strain also showed that TD+HR together resulted in a distribution of stresses and strains from the tongue (except for anterior) to below the hyoid, and that it also increased the stress and strain along the soft palate, constrictors and around the hyoid.

As for total tissue displacement, TD resulted in a displacement mainly of the thyroid, soft palate and the caudal end of the constrictors. HR, depending on the direction, resulted mostly in displacement of the tongue and the tissues around the hyoid bone. TD+HR resulted in the greatest tissue displacement with a combination of the aforementioned tissues (Figure 10).

### Caudal Baseline Hyoid Position Effects (OSA Phenotype)

Results in this section are for TD and anterior-cranial HR applied to the OSA-like phenotype of a more caudal baseline hyoid position.

#### Upper Airway Collapsibility (Pclose)

Compared to the original baseline hyoid position, Pclose was 23% and 34% higher for 2 and 4mm caudal baseline hyoid position (HR=TD=0 mm), respectively (Figure 11). TD alone at 3mm resulted in a decrease in Pclose of −3% for 2 mm caudal baseline hyoid position, and an increase in Pclose of 17% for 4 mm caudal baseline hyoid position; meanwhile, Pclose was −34% for the original baseline hyoid position. Anterior-cranial HR alone at 3mm resulted in a decrease in Pclose of −46% and −14% for 2 mm and 4 mm caudal baseline hyoid positions, respectively, while Pclose was −106% for the original baseline hyoid position. For TD+anterior-cranial HR, both at 3mm, the change in Pclose was −88% and −43% for 2 mm and 4 mm caudal baseline hyoid position, while Pclose was −131% for the original baseline hyoid position (Figure 11).

**Figure 11:**
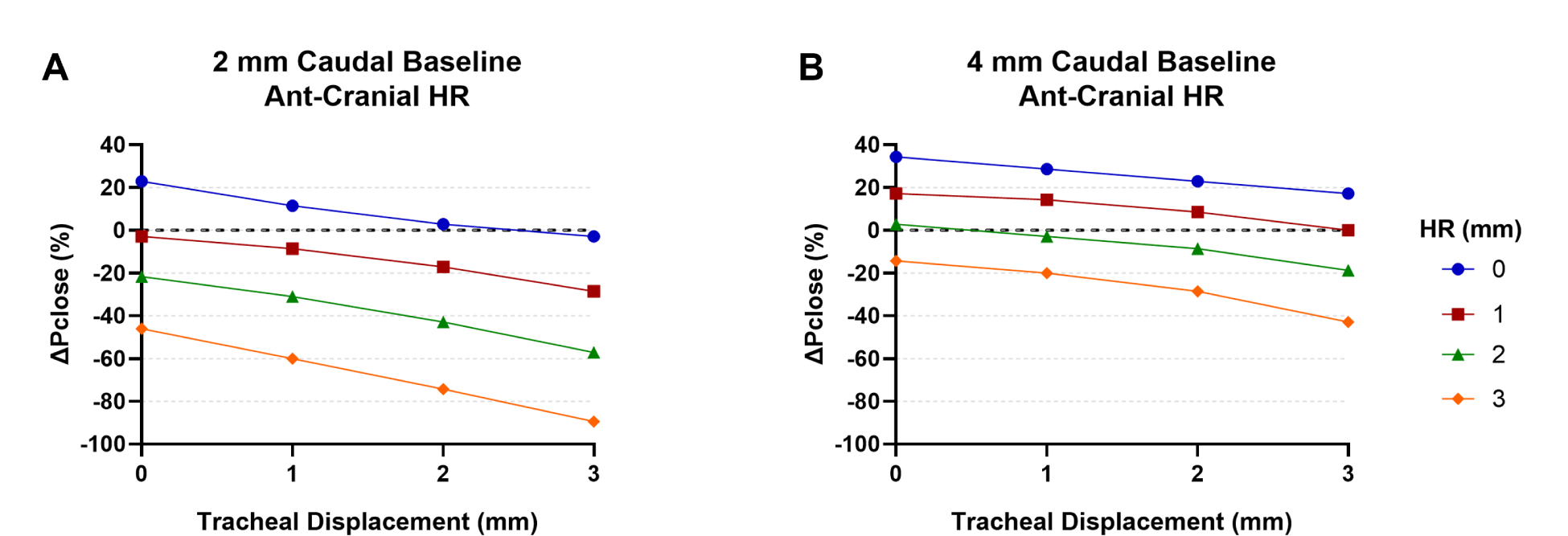
Pclose Outcomes for Caudal Baseline Hyoid Positions. Percent change in Pclose compared to the original baseline hyoid position for the 2 mm (A) and 4mm (B) caudal baseline hyoid positions with tracheal displacement (TD) and anterior-cranial (ant-cranial) hyoid repositioning (HR). Without HR and TD application (HR=TD=0mm; blue solid circle intersecting y-axis in A and B), Pclose was increased with a more caudal baseline hyoid position. Similar to the original baseline hyoid position, an increase in TD and ant-cranial HR alone resulted in a more negative Pclose, which decreased even further when combined. However, compared to the original baseline (Figure 5D), the more caudally positioned the hyoid, the less Pclose is reduced with TD and ant-cranial HR.

#### Upper Airway Geometry

##### Area

A more caudal baseline hyoid position did alter area compared to the original baseline hyoid position. TD alone at 3mm increased area by 19% and 18% for 2 mm and 4 mm caudal baseline hyoid positions, respectively, while the increase was 21% for the original baseline hyoid position. Anterior-cranial HR alone at 3 mm increased the area by 29% and 28% for 2 mm and 4 mm caudal baseline position, respectively, whereas the original baseline hyoid position area was increased by 32%. TD+anterior-cranial HR, both at 3 mm, resulted in an increase of 52% and 49% for 2 mm and 4 mm caudal baseline hyoid positions, respectively; meanwhile, the original baseline hyoid position resulted in a 55% increase due to both TD and HR together (Figure 12).

**Figure 12:**
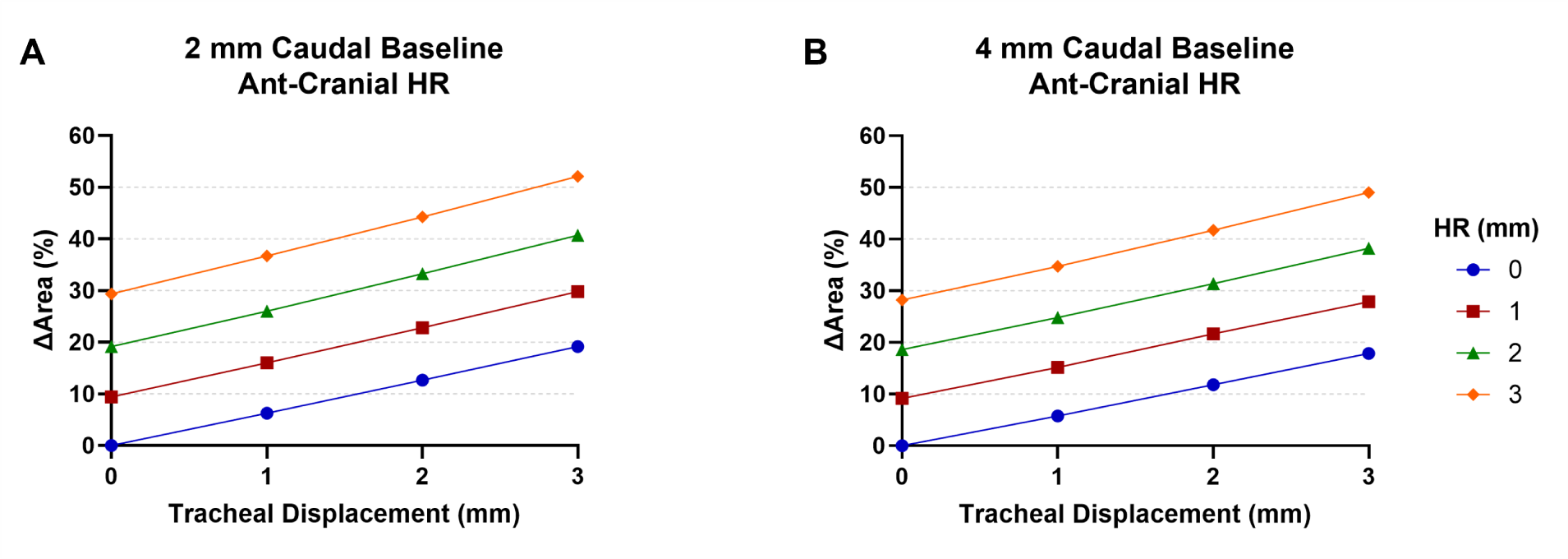
Upper Airway Area Outcomes for Caudal Baseline Hyoid Positions. The change in Area increased as tracheal displacement (TD) or hyoid repositioning (HR) were applied. TD+ant-cranial HR further increased area. The resulting increase in area for the 2 mm caudal baseline model at maximum displacement (52% increase in area at TD = 3 mm + ant-cranial HR = 3 mm) was slightly larger than that of the 4 mm caudal baseline (49% increase in area at TD = 3 mm and ant-cranial HR = 3 mm). The more caudally positioned the hyoid (4 mm vs. 2 mm), the lower the increase in area with TD+ant-cranial HR compared to the original baseline hyoid position (Figure 6D).

##### Anteroposterior Diameter

APD was not altered by a more caudal baseline hyoid position. TD alone increased APD in all regions, but mostly the R3 region at 3 mm (17% for both 2 mm and 4 mm caudal hyoid baseline positions). On the other hand, the original baseline hyoid position demonstrated the largest increase (18%) in the R2 region. Anterior-cranial HR alone resulted in a larger increase in all regions for a more caudal baseline hyoid position, most significantly for R2 at 3 mm (107% for 2 mm caudal hyoid and 102% for 4 mm caudal hyoid), which was similar for the original baseline hyoid position (largest increase of 107% in R2). TD+anterior-cranial HR at 3 mm for the caudal baseline hyoid positions resulted in larger increases in all regions than when either was applied alone, similar to the original baseline hyoid position, especially in R2 (123% for 2 mm caudal hyoid and 115% for 4 mm caudal hyoid, noting the change was 128% for the original baseline hyoid position) (Figure 13).

**Figure 13:**
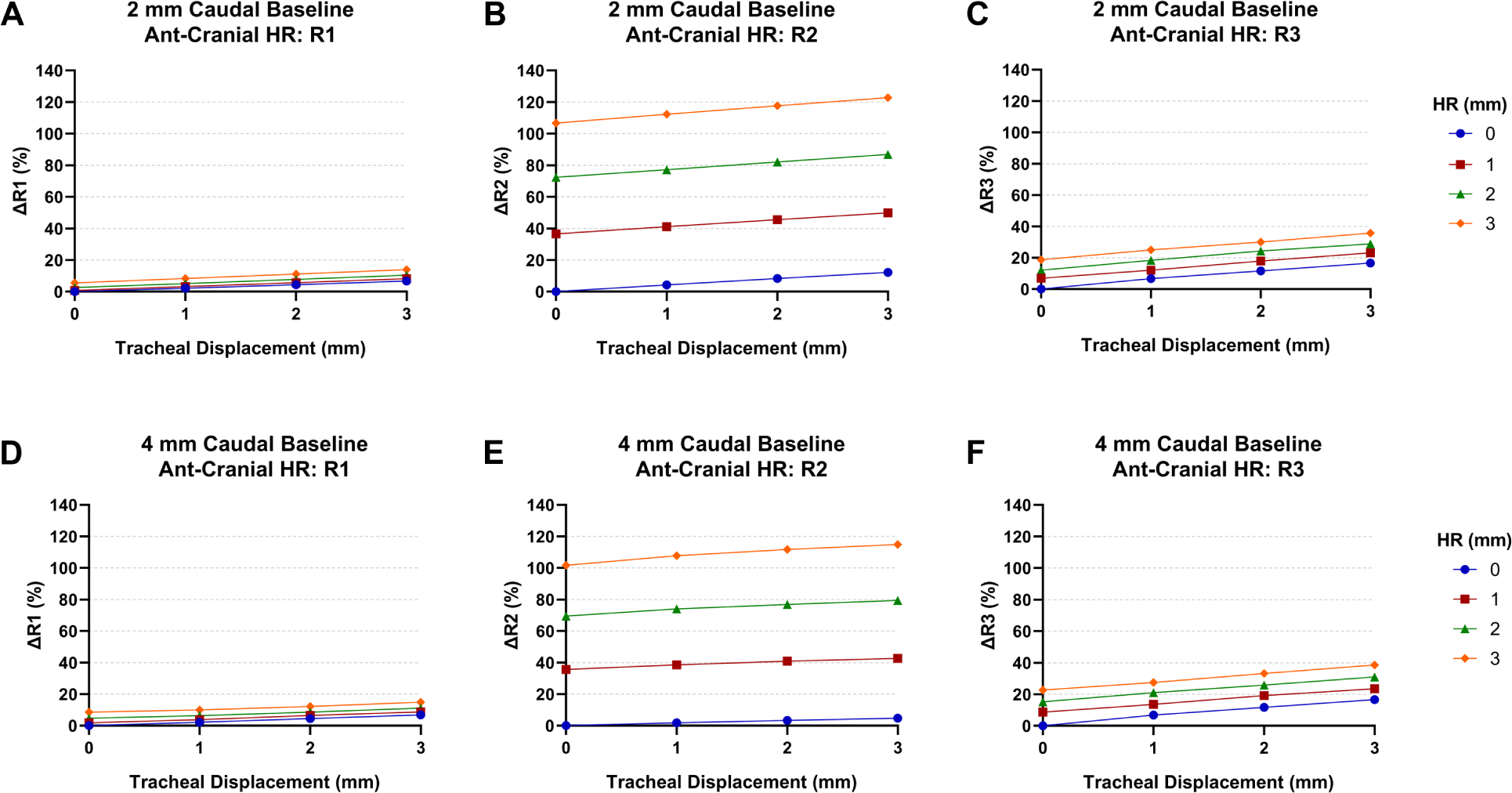
APD Outcomes for Caudal Baseline Hyoid Positions. Increasing tracheal displacement (TD) and anterior-cranial (ant-cranial) hyoid repositioning (HR), both separately and together resulted in an increase in anteroposterior diameter (APD) in all regions, especially R2, while R1 APD revealed the least change. A caudal baseline hyoid position of 2mm resulted in a larger increase in R2 APD compared to the 4 mm caudal baseline hyoid position. Thus, the more caudally positioned the hyoid, the less the increase in the APD around the hyoid in R2 (compare also with Figure 7 for the original baseline hyoid position).

#### Upper Airway Soft Tissue Mechanics (stress, strain and displacement)

TD+anterior-cranial HR resulted in the largest distribution of stress and strain particularly under the hyoid bone (Figures 14 & 15). A similar observation was noted for soft tissue displacement, which also indicated movements within regions with no direct displacement, such as the tongue (Figure 16). The patterns were similar between both caudal baseline hyoid positions compared to those of the original baseline hyoid position, however the more caudal the hyoid, the greater the resulting magnitudes on the soft tissues under the hyoid.

**Figure 14:**
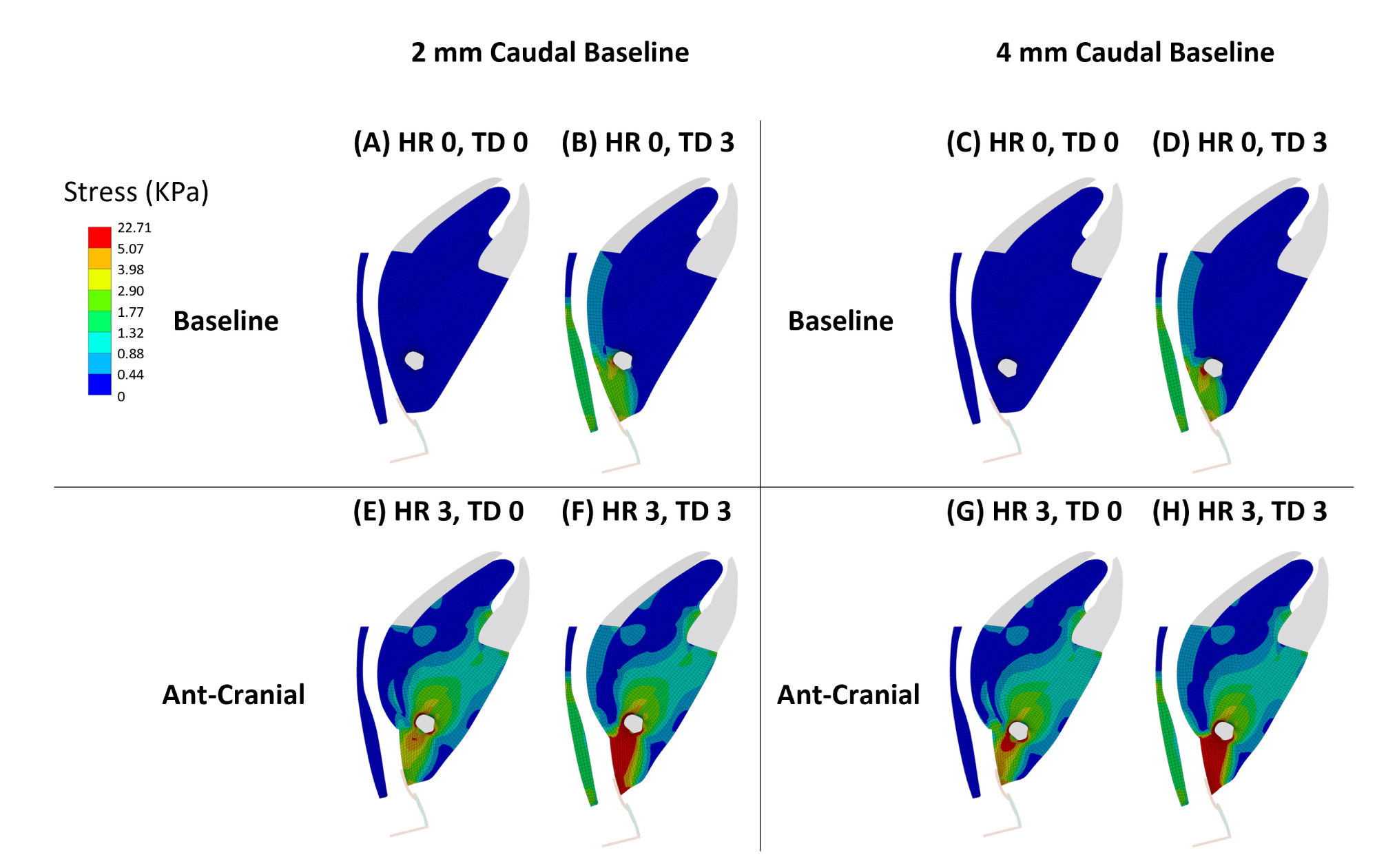
Soft Tissue Stress for the Caudal Baseline Hyoid Positions. Tracheal displacement (TD) and hyoid repositioning (HR) alone and in combination increased the distribution of stress in the soft tissues, yet the combination resulted in the greatest overall distribution. Stress distributions for 2 and 4 mm caudal baseline hyoid positions are similar to each other and those of the original hyoid baseline position (Figure 8). However, the resulting stress magnitudes are generally slightly higher for the 4 mm caudal baseline below the hyoid bone. HR 0 = HR at 0 mm. HR 3 = HR at 3 mm. TD 0 = TD at 0 mm. TD 3 = TD at 3mm.

**Figure 15:**
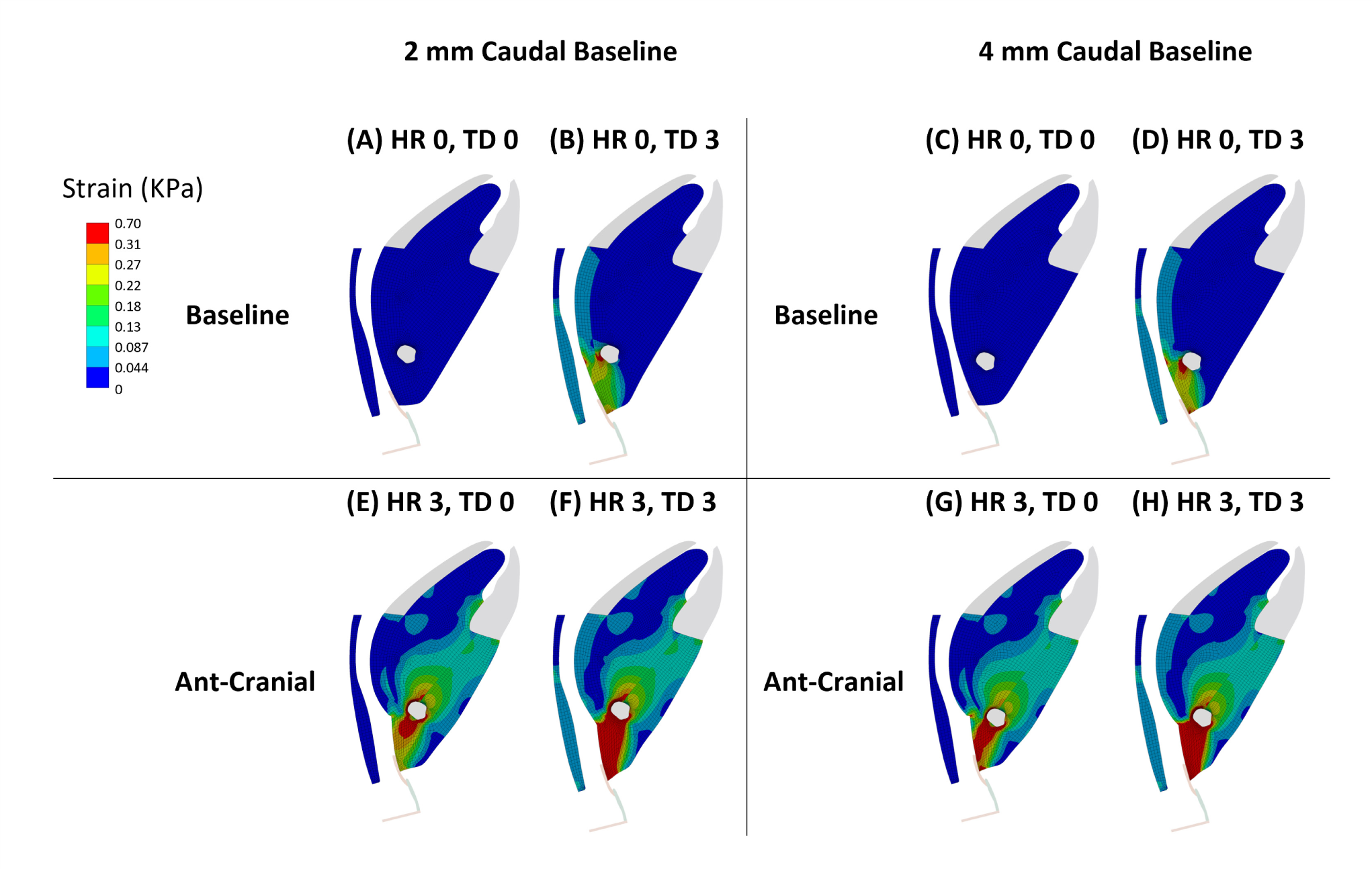
Soft Tissue Strain for the Caudal Baseline Hyoid Positions. Tracheal displacement (TD) and hyoid repositioning (HR) alone and in combination increased the distribution of strain in the soft tissues, yet the combination resulted in the greatest overall distribution. Strain distributions for 2 and 4 mm caudal baseline hyoid positions are similar to each other and those of the original hyoid baseline position. However, the resulting strains are generally slightly higher for the 4 mm caudal baseline under the hyoid bone. HR 0 = HR at 0 mm. HR 3 = HR at 3 mm. TD 0 = TD at 0 mm. TD 3 = TD at 3mm.

**Figure 16:**
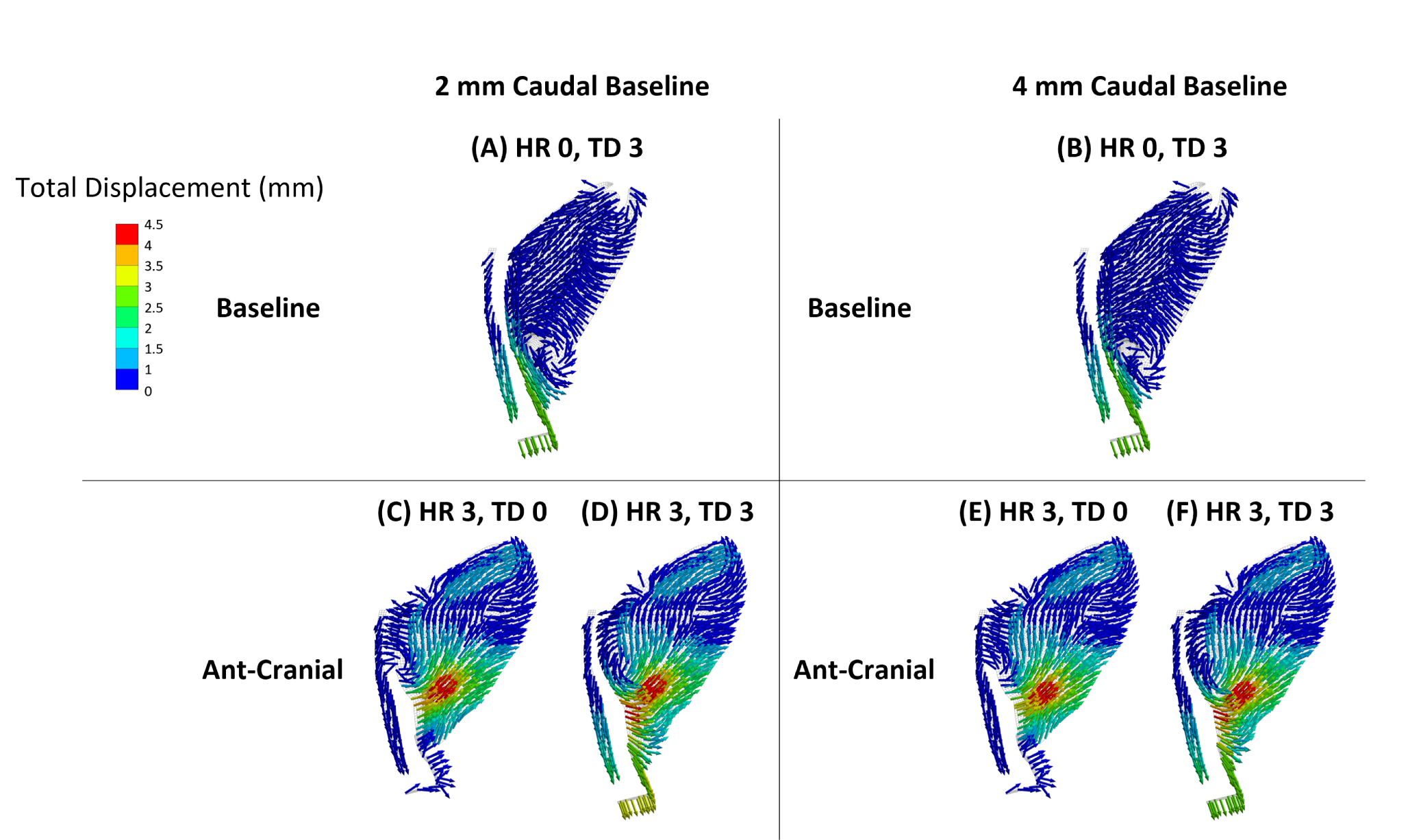
Tissue Displacement for the Caudal Baseline Hyoid Positions. Tissue displacements with a more caudal baseline hyoid position at 2 and 4 mm were similar to each other and the original baseline hyoid position (Figure 10). HR 0 = HR at 0 mm. HR 3 = HR at 3 mm. TD 0 = TD at 0 mm. TD 3 = TD at 3mm.

## Discussion

This finite element modeling study is the first to investigate the combined effects of TD and HR on upper airway outcomes related to collapsibility, geometry and tissue mechanics. Additionally, the influence of a more caudally positioned baseline hyoid bone, mimicking the OSA phenotype, on upper airway outcomes associated with TD and its combination with anterior-cranial HR, similar to hyomandibular suspension surgery, were also uniquely studied. Study findings revealed that the combination of both TD and HR (except in the caudal direction) decreased upper airway collapsibility and increased upper airway size, soft tissue stress and strain, more so than when either intervention was applied alone. A more caudally positioned baseline hyoid had negative effects on upper airway collapsibility, including reducing the overall positive impacts of TD and HR on all upper airway outcomes.

### Tracheal Displacement

TD is the primary mechanism by which lung volume affects upper airway patency and has been extensively studied (15–17, 25, 26). An increase in lung volume causes caudal TD, which has been shown to increase upper airway size, reduce collapsibility and stiffen upper airway soft tissues in animal, computational and physical models (14–16, 22, 27).

Similarly, in this study, increasing TD resulted in a gradual decrease in Pclose, increase in airway size, and an increase in stress and strain distributions and thus increased tissue stiffness. As a result, the upper airway became more stable with improved patency. TD also led to a nonuniform increase in upper airway size, particularly around the hyoid bone region (R2), consistent with previously reported outcomes (15, 16). In OSA, the pharyngeal muscles and tissues are thought to be softer than in healthy individuals (28–30). Therefore, TD, whether via static (e.g., body position, weight loss) or dynamic changes during breathing, likely plays a crucial role in stiffening the pharyngeal structure and is an important factor to consider for improving upper airway patency (25).

Note that unlike in our previous simulations of TD (16), the hyoid was fixed in the current study during load application. This approach was undertaken to maintain consistency with the HR simulations, where the hyoid was also fixed following load application, for direct comparisons. Indeed, this then allows for further analysis into the impact of hyoid movement on TD outcomes by comparing between the previous and current study. Since the hyoid was fixed in this study when applying TD, the resulting airway area and APD were slightly higher than those reported by Amatoury et al (16) for the same increment of 3mm, especially in the regions under the hyoid. Upper airway area due to TD alone in the current study was about 7% higher than those recorded by Amatoury et al (16). R1 APD was minimally greater by 0.5% in this study, but R2 and R3 (regions under the hyoid) APDs were about 9% higher than the previous study (16). The greater change in size/dimensions with TD in the current study compared to the previous study are due to the additional stretching of the tissues with TD when the hyoid is fixed, particularly below the hyoid bone, which resulted in an increase in size in these regions. However, fixing the hyoid bone means that TD loads are not effectively transferred to tissues above the hyoid, such as the tongue, which demonstrated reduced displacements, stress and strains compared to when the hyoid was free (16). However, stresses and strains are indeed greater below the hyoid with TD when the hyoid was fixed (16). Thus, with TD, a fixed hyoid (or at least with reduced mobility), may be beneficial to individuals with collapse in the lower airway region, while a hyoid free to move may be more beneficial to those with tongue-based collapse.

### Hyoid Repositioning

Improvements in the upper airway were dependent on the direction and increment of HR. Similar to our previous investigations, HR in anterior-based directions had the greatest impacts in reducing upper airway collapsibility and increasing upper airway size (4, 20). Additionally, in this study, we also demonstrated that dimension changes occur particularly in the hyoid region (R2). Furthermore, soft tissue stress, strain and displacement were also increased, especially around the hyoid and along the tongue.

Pclose was reduced following HR alone (except caudal HR), especially in the anterior-cranial direction. These findings align with previous research outcomes (4, 20). However, given the trachea in our model was fixed unlike in our previous study, we reported slightly greater changes in Pclose. This study noted a maximum decrease in Pclose of −26% for cranial HR and −106% for anterior-cranial HR, whereas Salman and Amatoury (20) reported a change of −20% for cranial HR and −96% for anterior-cranial HR (for the same increment of 3mm). The reason for the differences when the trachea was fixed with HR is similar in concept to when the hyoid was fixed with TD, as explained above for airway size. With the trachea fixed, there is greater stretch and stiffening of soft tissues with cranial-based HR directions, leading to further reductions in collapsibility.

As for airway geometry, HR alone independently increased upper airway area, except with caudal HR which decreased area. This decrease was a result of compression of the soft tissue mass under the bone with caudal HR, which narrowed the airway in the R2 and R3 regions, thus decreasing the overall area despite the increase in the R1 region. Conversely, anterior-based HR directions, particularly anterior-cranial, increased the area the most. Our previous research also revealed a similar trend in area with HR (20). However, again, due to the fixed trachea when applying HR in the current model, the increase in area in the cranial-based directions was 4% higher than that reported by Salman and Amatoury (20). The area outcomes for the rest of the HR directions in this study were in line with those previously presented by Salman and Amatoury (20).

Stress, strain and displacement of the soft tissues was increased when HR (in any direction) was applied. This increase was distributed across the soft tissues around the hyoid and tongue, which, depending on the HR direction, indicates the regions that may also be targeted and improved during surgical HR. The pattern of these outcomes is similar to the observations by Salman and Amatoury (20), but of relatively and slightly higher magnitudes because the trachea was fixed in this study.

### The Combination of Hyoid Repositioning and Tracheal Displacement

This study is the first to assess the combined effects of TD and HR. The combination of TD with HR (except in the caudal direction) augmented the improvement in upper airway outcomes compared to each intervention applied independently. TD+anterior-cranial HR induced the greatest improvement in airway size, collapsibility and tissue mechanics.

For Pclose, when both TD and HR were combined, there was a 25-97% improvement in Pclose depending on the direction of HR. For area, there was a 23-34% improvement when TD and HR were combined. The application of TD following HR (except caudal) stretched tissues even further, as indicated by the increased stress and strain. This resulted in a further decrease in collapsibility and increase in upper airway dimensions, especially around and under the hyoid in the R2 and R3 regions.

Previous research supports the notion that a combination of loads on the upper airway results in further improvements to the airway, especially when the loads are being applied in opposite direction. Rowley et al (26) demonstrated an interaction between passive tongue protrusion and TD that improved maximal flow and Pcrit outcomes in felines. Evidence of increased improvements in upper airway dimensions and stresses/strains has also been shown with combined mandibular advancement and TD in rabbit models (16, 31, 32). Furthermore, the combination of ansa cervicalis stimulation, which results in caudal TD, and hypoglossal nerve stimulation (HGNS) improves airflow by increasing the airway’s area even further than when HGNS is applied alone (33, 34). Thus, combined interventions, particularly with loads that act on the upper airway in different directions, seem to have the greatest impact on improving upper airway outcomes, as was the case in this study with TD+anterior-cranial HR.

### The Effect of a Caudal Hyoid Phenotype

The results of this study confirm that shifting the baseline hyoid position caudally, representing the OSA phenotype, negatively affects upper airway collapsibility, size and tissue mechanics following TD and HR interventions compared to the original baseline position. Thus, the baseline hyoid position is an important factor to consider in obtaining the desired upper airway outcomes with TD and HR interventions.

The caudal hyoid phenotype reduced the percentage of change in upper airway collapsibility and size for the same TD and HR increments and direction compared to the original baseline hyoid position. The more caudal the hyoid, the greater the negative influence, and this was most evident for Pclose. Thus, TD+HR interventions were less effective for the caudal hyoid phenotype. In fact, there is evidence that an increase in OSA severity, and hence lower hyoid position (35, 36), reduces the effectiveness of HR surgeries (37). Thus, consideration of baseline hyoid position, along with the degree of TD and direction/increment of HR, is likely required with HR surgeries for OSA for improved outcomes.

### Clinical Implications

The significance of the current study lies in its potential for clinical applications through two suggested strategies to improve upper airway functionality: personalized HR therapies or combination HR therapies.

Given the existence of different natural TD ranges (25), then outcomes of HR surgery may be further improved by optimizing surgery directions and increments based on the patient’s baseline hyoid position and natural range of TD during breathing and/or position changes while asleep.

On the other hand, a second strategy may be to combine HR surgery with stimulated TD, such as via ansa cervicalis nerve stimulation (33, 34), a promising new therapy for OSA. Indeed, in the current study, upper airway outcomes were further improved following both TD+HR, thus optimizing the increments of both interventions may improve the overall outcome especially as an increase in TD will aid in stiffening the upper airway further with HR, enhancing the outcomes of surgical therapies by improving airway stability and function. Additionally, the baseline hyoid position should be taken into consideration when undertaking HR surgeries and in combining them with TD to improve their effectiveness.

### Limitations and Critique of Methods

The general limitations of the model and interventions have been described in detail previously (16, 20), and here we limit our discussion to the current employed methodologies.

A rabbit model was chosen for its similar fundamental anatomical upper airway structure to humans, in addition to the consistency in upper airway outcomes to the human (4, 15–17, 20–23). In addition to its direct relevance to the human circumstance, utilizing a rabbit model allows continuous enhancement and validation from the experimental rabbit model from which it was originally built from and validated against for several interventions (4, 15, 16, 20, 24). Indeed, the original computational rabbit upper airway model (16) has also been used to validate a 2D model of the human upper airway by (38), further supporting the reliability of the current model.

Three TD and HR combinations (TD = 3 mm + anterior HR = 3 mm; TD = 2 mm + anterior-cranial HR = 3 mm; TD = 3 mm + anterior-cranial HR = 3 mm) applied to the model for measuring Pclose caused sizable deformation and distortion in the model that limited convergence, and so a Pclose value could not be directly obtained. This is because the repositioning magnitudes were relatively large, and coupled with the negative intraluminal pressure, caused extensive deformation on the model that exceeded its limits. Therefore, a linear regression model was used, based on the data available, to predict the Pclose results for the failed combinations. The linear regression model had an R^2^ of 0.9947, thus the outcomes are likely very close to what our finite element model would have predicted.

## Conclusion

This study has revealed that the combination of both tracheal displacement and hyoid repositioning (except the caudal direction) improved upper airway outcomes more than when either intervention was applied alone. The upper airway was less collapsible, stiffer and larger with both interventions in a magnitude dependent manner, with the greatest upper airway improvements occurring with anterior-cranial hyoid repositioning combined with tracheal displacement. Moreover, a more caudal baseline hyoid position, or OSA phenotype, reduced the degree of upper airway improvement with tracheal displacement and surgical hyoid positioning. Accordingly, this study provides proof of concept evidence of the importance of tracheal displacement for improved surgical hyoid repositioning therapy outcomes, or vice versa. It suggests the need to consider the baseline hyoid position, magnitude of tracheal displacement and magnitude/direction of surgical hyoid repositioning with such treatment interventions, which could lead to improved OSA treatment outcomes.

## Acknowledgements

The authors would like to thank Associate Professors Terence Amis and Kristina Kairaitis (The Westmead Institute for Medical Research and University of Sydney, Australia) and Professor Lynne Bilston (Neuroscience Research Australia and University of New South Wales, Australia) for their intellectual input and support.

## Author contributions

Model development and simulations were conducted at the American University of Beirut. JA conceived the research; DB and JA contributed to design of the research; DB performed the simulations; DB analyzed data; DB and JA interpreted the data; DB drafted the initial manuscript; DB and JA edited, revised, and approved final version of manuscript.

