## Supplementary Material for "Computational Simulations of Hyoid Bone Position and Tracheal Displacement: Effects on Upper Airway Patency and Tissue Mechanics"

This supplementary material includes figures that describe the model boundary conditions (Figure S1), contact conditions (Figure S2), and passive pharyngeal muscle connections (Figure S3). Model boundary conditions and contact conditions are indicated and labeled in Figures S1A and S2A, respectively, and their properties detailed in Figures S1B and S2B. Pharyngeal muscles, represented by linear elastic springs are shown in Figure S3A and details regarding stiffness properties are shown in Figure S3B.

Material properties of the model are described in Tables S1 and S2 for linear elastic materials and non-linear hyperelastic parameters, respectively.

### Supplementary Figures

A

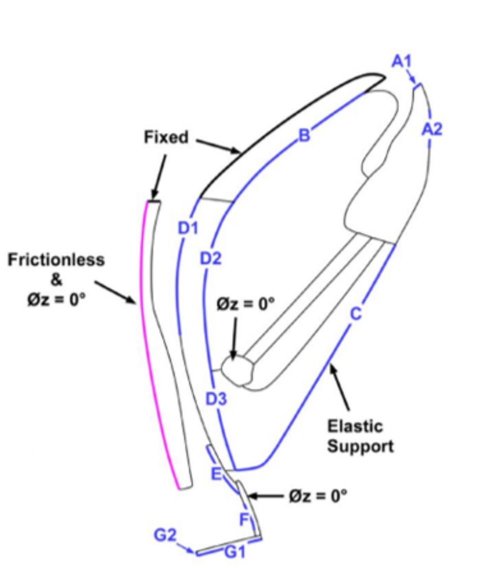


B

| Tissue | Boundary Label | Elastic Foundation Stiffness, mN/mm$^2$ |
|---|---|---|
| **Mandible** | A1 | 2,000 |
| | A2 | |
| **Tongue** | B | 1 |
| **Tissue Mass** | C | 0.5 |
| **Soft Palate** | D1 | 0.8 |
| | D2 | 0.1 |
| | D3 | 0.1 |
| **Epiglottis** | E | 5 |
| **Thyroid Cartilage** | F | 50 |
| **Base plate** | G1 | 100 |
| | G2 | 50 |

**Figure S1: Model boundary conditions**. (A) Illustration of the model and several boundary conditions indicated in different colors: blue represents elastic supports (labels A1 to G1), black represents fixed supports, pink represents frictionless supports and no rotation condition (Øz = 0°) is applied to the constrictors, thyroid cartilage and hyoid bone. (B) Elastic constraint properties corresponding to each of the boundaries labeled in (A). Figure adapted from Salman and Amatoury (1).

A

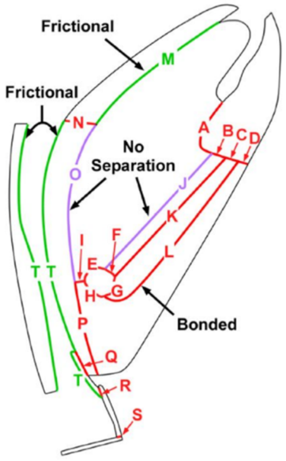


B

| Interface | Label | Behavior | Stiffness Factor |
|---|---|---|---|
| **Mandible/attached tissues** | A-D | Bonded | 1 |
| **Hyoid bone/attached tissues** | E and G-H | Bonded | 1 |
| **Hyoid bone/geniohyoid** | F | Bonded | 2 |
| **Tongue/tissue mass** | I | Bonded | 1 |
| **Tongue/geniohyoid** | J | No separation | 1 |
| **Geniohyoid/mylohyoid** | K | Bonded | 1 |
| **Mylohyoid/tissue mass** | L | Bonded | 1 |
| **Tongue/hard palate** | M | Frictional, $\mu = 0.2$ | 1 |
| **Hard palate/soft palate** | N | Bonded | 1 |
| **Soft palate/tongue** | O | No separation | 1.55 |
| **Soft palate/tissue mass** | P | Bonded | 1 |
| **Soft palate/epiglottis** | Q | Bonded | 1 |
| **Thyroid cartilage/epiglottis** | R | Bonded | 1 |
| **Thyroid cartilage/base plate** | S | Bonded | 1 |
| **Soft palate-epiglottis/constrictors** | T | Frictional, $\mu = 0.1$ | 1 |

**Figure S2: Model contact conditions** (A) Illustration of the model and several contact conditions indicated in different colors: red indicates bonded contacts, purple indicates no separation and green indicates frictional contacts. (B) Model contact interfaces labeled in (A) and their respective properties such as contact condition behavior and normal stiffness factor. Figure adapted from Salaman and Amatoury (1).

**A**

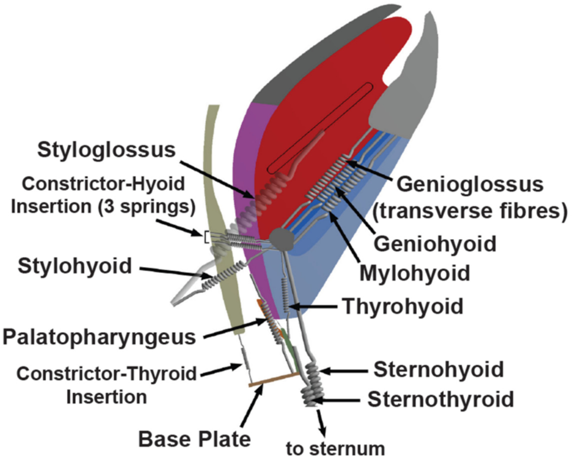


**B**

| Spring (Muscle) | Stiffness, N/mm |
|---|---|
| **Genioglossus (transverse fibers)** | 0.3 |
| **Geniohyoid** | 0.15 |
| **Mylohyoid** | 0.15 |
| **Styloglossus** | 0.001 |
| **Stylohyoid** | 0.6 |
| **Thyrohyoid** | 0.25 |
| **Sternothyroid** | 0.2 |
| **Sternohyoid** | 0.2 |
| **Soft Palate-base plate** | 0.0005 |
| **Constrictors-hyoid (1)** | 0.1 |
| **Constrictors-hyoid (2)** | 0.1 |
| **Constrictors-hyoid (3)** | 0.5 |
| **Constrictors-base plate** | 0.008 |
| **Soft palate-thyroid** | 0.00001 |
| **Tissue mass-thyroid** | 0.005 |

**Figure S3: Passive pharyngeal muscle connections**. (A) Passive action of several pharyngeal muscles of the rabbit upper airway modeled are modeled as linear elastic springs. (B) Stiffness properties corresponding to each of the springs representing different pharyngeal muscles. Figure adapted from Salman and Amatoury (1).

### Supplementary Tables

**Table S1:** Linear elastic material for bony and cartilaginous tissues

| Tissue | Parameters | |
|---|---|---|
| | **E (MPa)** | $\upsilon$ |
| **Hyoid Bone**<br>**Mandible**<br>**Hard Palate**<br>**Base Plate**<br>**Thyroid Cartilage** | 17000 | 0.49 |
| **Epiglottis** | 2.4 | 0.49 |

E, Young's Modulus. $\upsilon$, Poisons ratio. Table adapted from Salman and Amatoury (1).

**Table S2:** Nonlinear hyperelastic model parameters (Yeoh second-order) for soft tissues

| Tissue | $\upsilon$ | $C_{10}$ (Pa) | $C_{20}$ (Pa) | $d_{10}$ ($Pa^{-1}$) $\times 10^{-5}$ | $d_{20}$ ($Pa^{-1}$) $\times 10^{-5}$ |
|---|---|---|---|---|---|
| **Tongue**<br>**Soft Palate** | 0.49 | 1628 | 770 | 1.24 | 2.61 |
| **Tissue Mass**<br>**Geniohyoid**<br>**Mylohyoid** | 0.45 | 1532 | 725 | 6.75 | 14.3 |
| **Constrictor Body** | 0.45 | 4005 | 1896 | 2.58 | 5.46 |

Parameters are for the Yeoh second-order strain energy function given by, $W = C_{10}(I_1 - 3) + C_{20}(I_1 - 3)^2 + \frac{1}{d_{10}}(J - 1)^2 + \frac{1}{d_{20}}(J - 1)^4$. Table adapted from Salman and Amatoury (1).
